# SOX4 facilitates PGR protein stability and FOXO1 expression conducive for human endometrial decidualization

**DOI:** 10.1101/2021.07.19.452960

**Authors:** Pinxiu Huang, Wenbo Deng, Haili Bao, Zhong Lin, Mengying Liu, Jinxiang Wu, Xiaobo Zhou, Manting Qiao, Yihua Yang, Han Cai, Faiza Rao, Jingsi Chen, Dunjin Chen, Jinhua Lu, Haibin Wang, Aiping Qin, Shuangbo Kong

## Abstract

The establishment of receptive endometrium in human necessitates appropriate decidualization of stromal cells, which involves steroids regulated periodic transformation of endometrial stromal cells during menstrual cycle. Insufficient decidualization of endometrium contributes to not only the failure of embryo implantation and unexplained infertility, but also the occurrence of recurrent spontaneous abortion, intrauterine growth retardation, preeclampsia, and other clinical gynecological diseases. However, the potential molecular regulatory mechanism underlying the initiation and maintenance of decidualization in humans is yet to be fully elucidated. In this investigation, we document that SOX4 is a key regulator of human endometrial stromal cells (hESCs) decidualization by directly regulating PRL and FOXO1 expression as revealed by whole genomic binding of SOX4 assay and RNA-Seq. Besides, our immunoprecipitation and mass spectrometry results unravel that SOX4 modulates progesterone receptor (PGR) stability through repressing E3 ubiquitin ligase HERC4 mediated degradation. More importantly, we provide evidence that dysregulated SOX4-HERC4-PGR axis is a potential cause of defective decidualization and recurrent implantation failure (RIF) in IVF patients. In summary, this study evidences that SOX4 is a new and critical regulator for human endometrial decidualization, and provides insightful information for the pathology of decidualization-related infertility and will pave the way for pregnancy improvement.

## Introduction

Adequate crosstalk between implantation-competent embryo and receptive endometrium is prerequisite for successful pregnancy(Cha, Sun et al. 2012). The receptive endometrium in human requires remodeling of stromal cells under the regulation of rising progesterone and intracellular cyclic AMP, which will undergo more extensive transformation driven by factors secreted from embryos. This transformation renders necessary nutrition, immune tolerance, and resistance to oxidative stress for the development of implanted embryos and directs the invasion of embryonic trophoblasts(Duc-Goiran, Mignot et al. 1999, Fujiwara, Ono et al. 2020). Insufficient decidualization in endometrium is related to failed embryo implantation, unexplained infertility, recurrent spontaneous abortion(Coulam 2016), intrauterine growth retardation(Lefevre, Palin et al. 2011), and preeclampsia(Garrido-Gomez, Dominguez et al. 2017). However, the underlying molecular mechanism governing the endometrial decidualization remains enigmatic(Okada, Tsuzuki et al. 2018).

A wide range of transcription factors crucial for stromal cell decidualization have been successively identified(Gellersen and Brosens 2014). FOXO1, a member of FOXO fork-head transcription factors subfamily, is one of the earliest identified transcriptional factors in human endometrial stromal cells (hESCs) responding to decidualization stimulation (progesterone and cAMP)(Labied, Kajihara et al. 2006). Accumulative evidence has demonstrated that FOXO1 regulates transcription of PRL and IGFBP1 through direct binding to their promoters(Christian, Zhang et al. 2002, Kim, Buzzio et al. 2005). Progesterone receptor (PGR), which is a critical master factors for the endometrial stromal cells (ESCs) decidualization, imparts endometrium receptivity through binding with P4 and its nuclear translocation(Keller, Wiest et al. 1979, Mulac-Jericevic, Mullinax et al. 2000). An array of PGR direct target genes had been unraveled by PGR ChIP-Seq in both human and mice(Rubel, Lanz et al. 2012, Mazur, Vasquez et al. 2015, Chi, Wang et al. 2020). Abnormal PGR expression is closely relevant with unexplained infertility(Keller, Wiest et al. 1979) and endometriosis(Zhou, Fu et al. 2016, Pei, Liu et al. 2018). Although many transcription factors and downstream events in decidualization of mice and human have been unraveled(DeMayo and Lydon 2020), the precise mechanism orchestrating transcriptional regulatory network underpinning endometrial decidualization remained not fully explored.

SOX4 is a highly conserved transcription factor belonging to the SOX (SRY-box) family. Studies have shown that SOX4 is vital to a variety of biological processes, including embryogenesis, neural development, and differentiation(Moreno 2020). Further, it has been noticed that SOX4 knockout mice die of cardiac malformation on the fourteenth day of pregnancy, suggesting the key role of SOX4 in embryonic development(Ya, Schilham et al. 1998). In addition, an increasing number of reports indicate that SOX4 is related to tumor cell proliferation, metastasis, and epithelial-mesenchymal transformation(Li, Liu et al. 2020). Report also shows that SOX4 is highly expressed in breast cancer under the regulation of progesterone(Graham, Hunt et al. 1999). Additionally, SOX17, another member of the SOX family, has also been observed to be a direct target of PGR in epithelium regulating IHH expression(Wang, Li et al. 2018). While the significance and regulation of SOX4 in female pregnancy remain intangible.

Here, we provide evidence that SOX4, under the regulation of P4-PGR, guides human endometrium stromal cells (hESCs) decidualization by regulating PRL and FOXO1 expression as revealed by ChIP-Seq and RNA-Seq. Mechanism studies also unravel the importance of SOX4 in maintaining the protein stability of PGR by repressing ubiquitin E3 ligase HERC4. Moreover, both SOX4 and PGR have been demonstrated to be aberrantly downregulated in the endometrium of endometriosis (EMS) patients suffering from implantation failure.

## Results

### SOX4 is dynamically expressed in hESCs regulated by P4-PGR signaling

To investigate the expression of transcription factors in hESCs, RNA sequencing (RNA-Seq) was performed in normal endometrium stromal cells. We noted that SOX4 was the 17^th^ highest expressed transcript factor with CCAAT enhancer-binding proteins (CEBPB, CEBPD) and ATF4 (Activating transcription factor 4) being the most abundant transcription factors in endometrium stromal cells (Fig. 1A, B). Meanwhile, previous RNA-Seq data in isolated stromal and epithelial cells revealed that SOX4 expression was the most abundant SOX family in stromal cells (Fig. S1A), while the expression of SOX17 was restricted to the epithelium(Deng, Yuan et al. 2019), which was consistent with the indispensable role of SOX17 for embryo implantation by regulating epithelial IHH expression(Wang, Li et al. 2018). Thus, we speculated that the conserved expression of SOX4 in stroma was linked to decidualization. To explore whether SOX4 was under the regulation of dynamic change of estrogen and progesterone during the menstrual cycle, we first analyzed the expression of SOX4 in endometrial biopsy samples obtained from healthy, reproductive-aged volunteers with regular menstrual cycles. Albeit SOX4 was detected at a low level in the E2-dominant proliferative phase, its expression was progressively increased in the nucleus of stromal cells in P4-dominant early, middle, and late secretory phases (Fig. 1C).

**Fig. 1.**
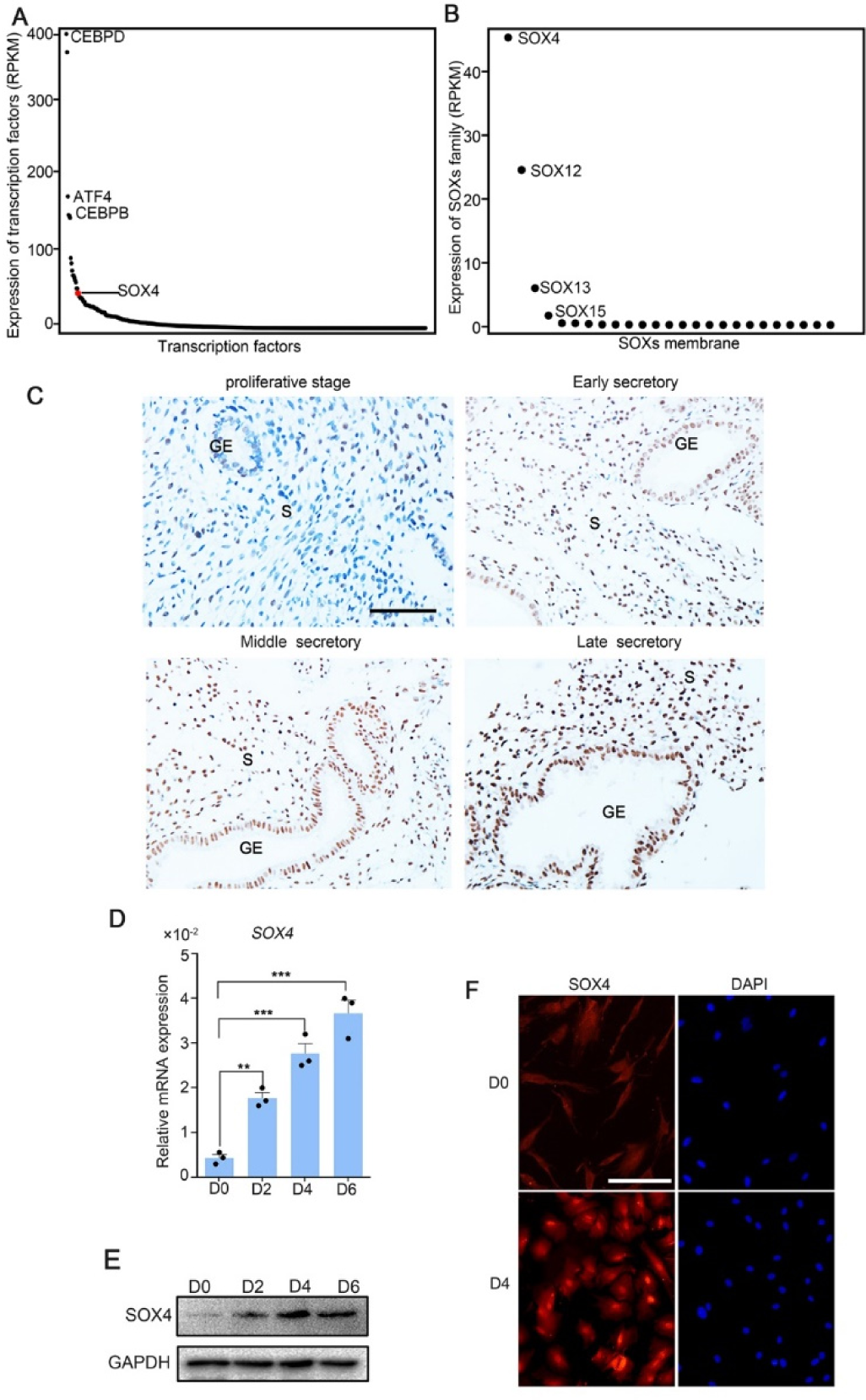
SOX4 is dynamically expressed in human ESCs. (A) Expression of all transcription factors in human non-decidualized ESCs by RNA-Seq. (B) Expression of SOX family genes in human non-decidualized ESCs by RNA-Seq. (C) Immunohistochemical analysis of SOX4 protein expression in proliferative, and secretory phases (early, middle, and late) of the menstrual cycle. GE: Gland epithelium; S: stromal. Scale bar: 100 μm. (D) Expression of *SOX4* mRNA levels in decidualized stromal cells at different time points. Results are presented as means ± SEM; n = 3; ** indicates P<0.005; *** indicates P<0.0001. (E) Expression of SOX4 protein levels in decidualized stromal cells at different time points. (F) Immunofluorescent detection of SOX4 protein localization in the undecidualized and decidualized hESCs. Scale bar: 100 μm.

This upregulation of SOX4 in differentiated stromal cell in secretory phase was also substantiated in *in vitro* decidualized stromal cells induced by E2, MPA and cAMP (EPC) cocktail with gradually cumulated mRNA and protein levels in immortalized hESCs (Fig. 1D, E) as well as in primary stromal cells (Fig. S2A, S2B). Immunofluorescence also manifested that SOX4 was mainly localized in the nucleus of hESCs after EPC treatment (Fig. 1F).

The intense expression of SOX4 in the stroma of secretory phase accompanied with rising P4 and inducted by EPC in in vitro stromal cells, suggesting that progesterone may be involved in the regulation of SOX4 expression. This assumption was supported by the observation that SOX4 expression was induced by MPA, a progesterone analogue commonly used in decidualization induction, and abrogated in the presence of PGR antagonist RU486 (Fig. 2A). Progesterone induced SOX4 mRNA and protein levels were significantly attenuated with PGR knockdown in immortalized hESCs (Fig. 2B-D). Thus, the P4-PGR signaling was critical for SOX4 expression in hESCs. Based on previous ATAC-Seq data from undifferentiation and differentiation stromal cells and PGR ChIP-Seq from proliferative and secretory endometrium(Chi, Wang et al. 2020), we found that there was intensive potential PGR binding at SOX4 promoter (Fig. 2E). This speculation was validated by PGR ChIP-qPCR at SOX4 promoter in decidualized stromal cells (Fig. 2F). Furthermore, the binding of PGR on SOX4 promoter was also confirmed by luciferase reporter assay that overexpression of PGR increased the reporter activities in the presence of MPA in 293T cells (Fig. 2G). These results indicated that PGR guided SOX4 expression via direct transcriptional regulation.

**Fig. 2.**
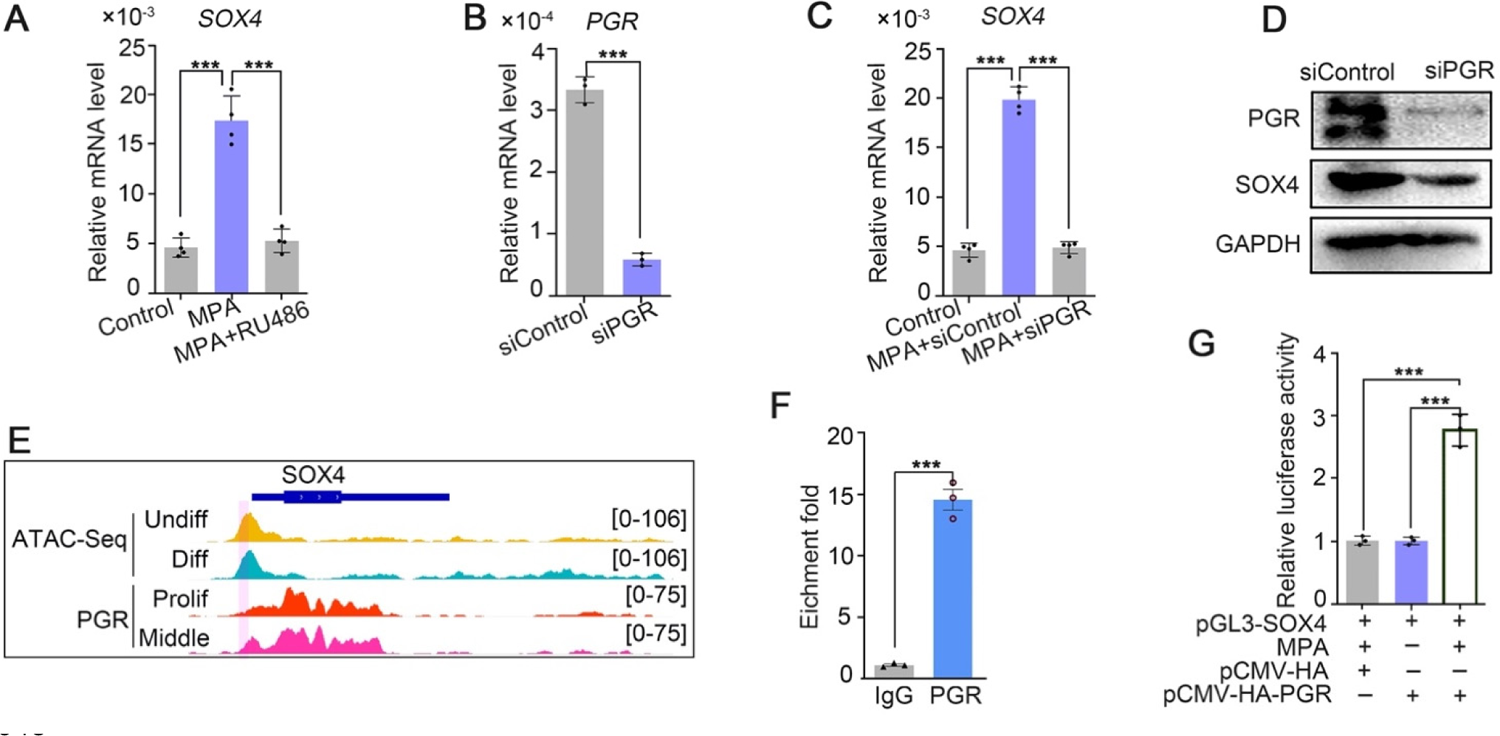
SOX4 is regulated by P4-PGR signaling in human ESCs. (A) Expression of *SOX4* mRNA in the presence of MPA or MPA+RU486 in immortalized hESCs. Results are presented as means ± SEM; n = 3; *** indicates P<0.0001. (B) Knockdown efficiency of PGR by siPGR. Results are presented as means ± SEM; n = 3; *** indicates P<0.0001. (C) Expression of *SOX4* mRNA in the presence of MPA with PGR knockdown. Results were presented as means ± SEM; n = 3; *** indicates P<0.0001. (D) Expression of SOX4 and PGR after PGR knockdown. (E) Visualization of PGR binding and chromatin accessibility on SOX4 locus. The chromatin accessibility is depicted in undifferentiated and differentiated stromal cells and genome-wide PGR binding is generated from proliferated and middle secretory endometrium as revealed from previous reports. Undiff: undifferentiated hESC; Diff: differentiated hESC; Prolif: proliferative endometrium; Middle: middle phase of secretory endometrium. (F) ChIP assay of potential PGR binding on SOX4 as indicated from (E) in decidualized immortalized hESC for 2 days. Data are plotted as mean ± SEM; n = 3; *** indicates P<0.0001.(G) Luciferase activity assay of SOX4 in the presence of MPA and PGR in 293T cells. Results are presented as means ± SEM. n = 3; * indicates P<0.05; ** indicates P<0.005; *** indicates P<0.0001.

### SOX4 is required for hESCs decidualization

To depict the significance of SOX4 during decidualization, SOX4 was knockout by CRISPR/Cas9 which was ascertained by Sanger-sequencing in both alleles (Fig. S2C) and further verified by western-blot and immunofluorescence (Fig. 3A and Fig. S2D). The expression of PRL and IGFBP1 were dramatically decreased in SOX4 knockout hESCs after decidualization for 2, 4, 6 days compared with SOX4 intact cells (Fig. 3B-D). Similar results were obtained in primary hESC after SOX4 knockdown by shRNA prior to EPC administration (Fig. S2E-H). On the other side, overexpressing SOX4 in hESCs significantly augmented the expression of *PRL* and *IGFBP1* in both immortalized and primary hESCs decidualized for four days (Fig. 3E-G and Fig. S2I-K).

**Fig. 3.**
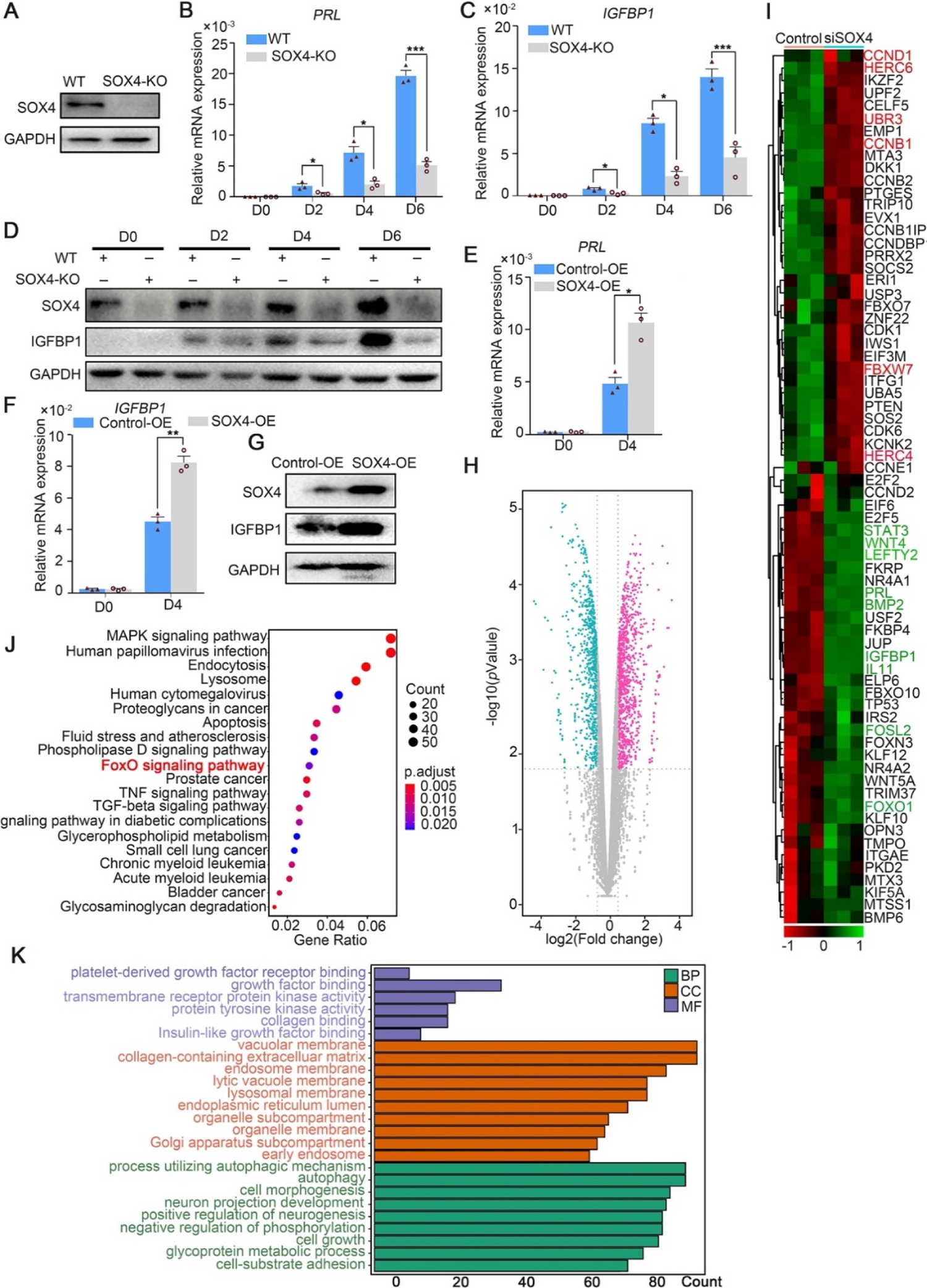
SOX4 regulated genes in decidualized hESCs. (A) Western blot verification of SOX4 knockout efficiency by sgSOX4. (B, C) mRNA levels of *PRL* (B) and *IGFBP1* (C) in undecidualized and decidualized immortalized hESCs after SOX4 knockout at indicated time points. Results are presented as means ± SEM; n = 3; * indicates P<0.05; *** indicates P<0.0001. (D) Protein levels of IGFBP1 in undecidualized and decidualized hESCs after SOX4 knockout at indicated time points. (E and F) mRNA levels of *PRL* (E) and *IGFBP1* (F) after SOX4 overexpression in undecidualized or decidualized hESCs for 4 days. n = 3; * indicates P<0.05.G) Protein levels of SOX4 and IGFBP1 in decidualized hESC for 4 days with control or SOX4 overexpression. (H) Differential expressed genes detected by RNA-Seq in immortalized hESCs decidualized for 2 days with control or SOX4 knockdown as visualized by volcano plot. (I) Heatmap of top differential expressed genes from RNA-Seq after SOX4 knockdown in decidualized hESCs. (J and K) KEGG (J) and GO (K) analysis of the differentially expressed genes in RNA-Seq. Results were presented as means ± SEM. * indicates P<0.05; ** indicates P<0.005; *** indicates P<0.0001. The above experiments were repeated three times.

Moreover, RNA-Seq was utilized in SOX4 intact or depleted immortalized hESCs treated with EPC for two days. The genes significantly downregulated after SOX4 knockdown included aforementioned *IGFBP1* and *PRL* as well as other genes critical for decidualization, such as *FOXO1*, *LEFTY2*, *WNT5A*, *WNT4*, *FOSL2*, *STAT3*, *LEFTY2*, *IL-11*, *BMP2* (Fig. 3H, I). Kyoto Encyclopedia of Genes and Genomes (KEGG) analysis showed that the differentially expressed genes were enriched in the FOXO signaling pathway, TGF-beta signaling pathway, and MAPK signaling pathway (Fig. 3J), consistent with the observed defective decidualization in the absence of SOX4. The Gene Ontology (GO) annotation analysis showed that the differentially expressed genes were related to autophagy-associated pathway, insulin-like growth factor binding, collagen-containing extracellular matrix, cell-substrate adhesion (Fig. 3K). Together, these results revealed the critical role of SOX4 in hESCs decidualization.

### Genome binding of SOX4 in decidualized stromal cells

To interrogate genes directly regulated by SOX4, we detected the chromatin-wide binding of SOX4 by ChIP-Seq. Our results showed that there were total of 23,709 SOX4 binding peaks with most of them enriched in promoters, distal intergenic and intron (Fig. 4A). Peak enrichment indicated that SOX4 binding was primary closed to TSS as well as evidenced by heatmap of peak distribution (Fig. 4B, C). We also noticed that SOX4 mainly binds to those genes with higher expression (RPKM >= 1) accompanied with few SOX4 binding on those less to no expression genes (RPKM < 1) (Fig. 4B). Motif analysis results revealed that the most significantly enriched motif was SOX4 consensus binding sites, further supported the reliability of this SOX4 ChIP-Seq. We also noticed that FRA2 and FOSL2, two critical members of AP1, were also highly enriched at SOX4 binding sites, indicating the potential cooperation between SOX4 and AP1 (Fig. 4D). Considering the positive correlation between SOX4 binding and gene expression, we overlapped SOX4 binding genes with those down-regulated genes after SOX4 knockdown and found that 389 genes with SOX4 binding decreased after SOX4 ablation, indicating the direct regulation of SOX4 on these genes (Fig. 4E).

**Fig. 4.**
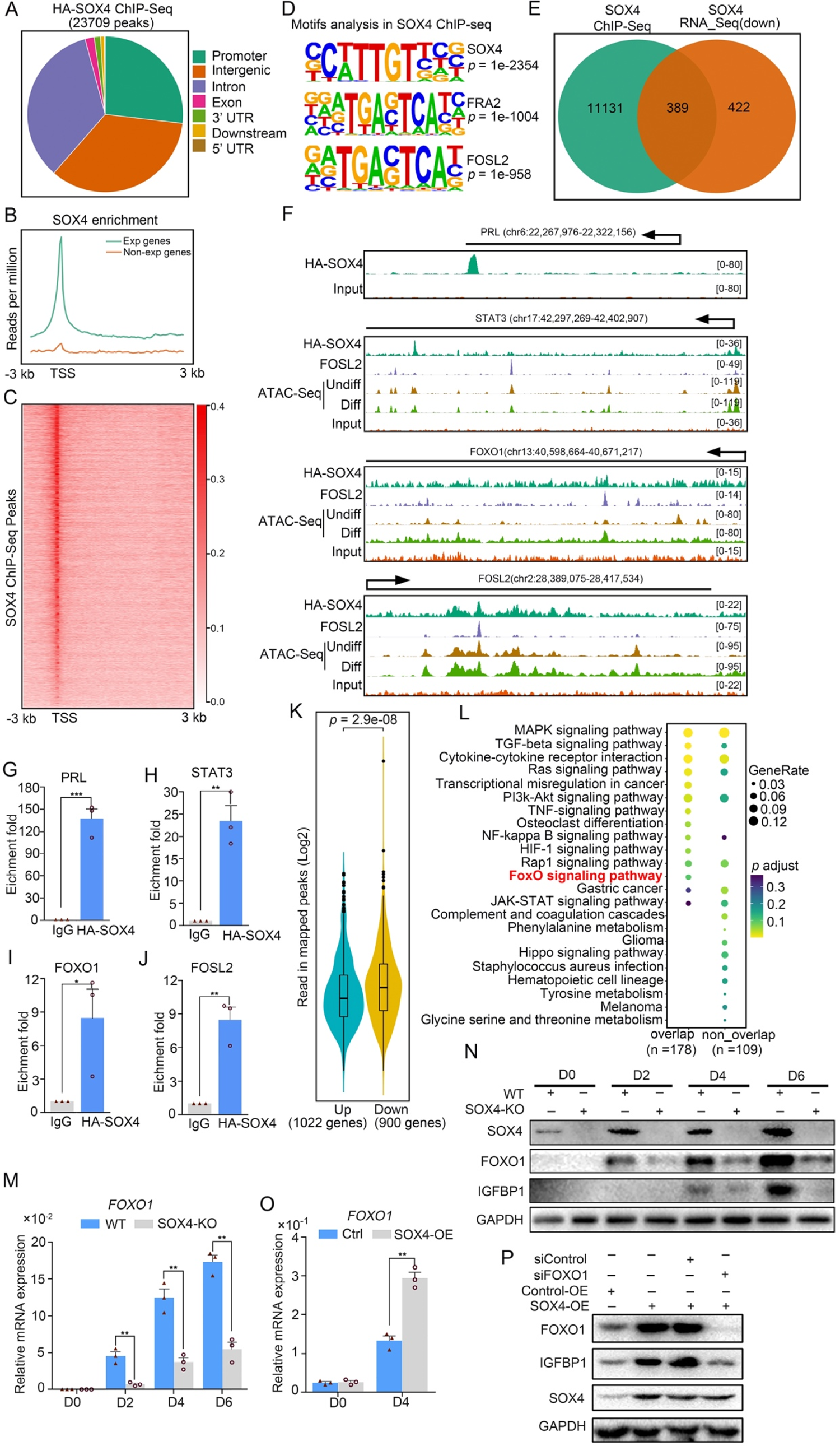
Genome wide binding of SOX4 in decidualized stromal cells. (A) Distribution of SOX4 binding peaks as revealed from SOX4 ChIP-seq in SOX4 overexpressed (HA-SOX4) immortalized hESCs decidualized for 2 days. (B) Distribution SOX4 binding in genebody. (C) Heatmap of SOX4 binding sites distribution. (D) Motif analysis of SOX4 binding sites. (E) Venn diagram of SOX4 directly binding peaks and SOX4 regulated genes after SOX4 knockdown. (F) SOX4 binding site in the PRL, STAT3, FOXO1, FOSL2 with PGR binding and chromatin accessibility. The chromatin accessibility is depicted in undifferentiated and differentiated stromal cells and genome-wide PGR binding is generated from proliferated and middle secretory endometrium as revealed from previous reports. Undiff: undifferentiated hESCs; Diff: differentiated hESCs; Prolif: proliferative endometrium; Middle: middle phase of secretory endometrium. (G-J) ChIP-qPCR assay of SOX4 binding on PRL, STAT3, FOXO1 and FOSL2 in HA-SOX4 overexpressed decidualized hESCs. Results were presented as means ± SEM; n=3; * indicates P<0.05; ** indicates P<0.005; *** indicates P<0.001. (K) Read number of SOX4 binding in SOX4 regulated genes after SOX4 knockdown. (L) KEGG analysis of overlapped genes of SOX4 directly binding and SOX4 downregulated genes as well as non-overlapped genes. (M) mRNA levels of *FOXO1* in SOX4 knockout undecidualized and decidualized hESCs at indicated time points. Results were presented as means ± SEM; n=3; **indicates P<0.005. (N) Protein levels of SOX4, FOXO1 and IGFBP1 in SOX4 knockout undecidualized and decidualized hESCs at indicated time points. (O) mRNA levels of *FOXO1* after SOX4 overexpression in undecidualized or decidualized hESCs for 4 days. Results were presented as means ± SEM; n =3; **indicates P<0.005. (P) Protein levels of FOXO1, IGFBP1 and SOX4 after overexpression of SOX4 in the presence of FOXO1 knockdown or not in decidualized hESCs for four days. The above experiments were repeated three times.

It was gratifying that there were several latent SOX4 binding sites on the PRL, STAT3, FOXO1 and FOSL2 (Fig. 4F). ChIP-qPCR also confirmed the binding of SOX4 on these genes (Fig. 4G-J). Those down-regulated genes (900) after SOX4 knockdown possess more SOX4 binding reads compared with upregulated genes (1,022) (Fig. 4K), indicating the transcriptional activation effect of SOX4. Importantly, those SOX4 direct target genes also showed specific enrichment of the FOXO signaling pathway (Fig. 4L). To further assess the regulation of SOX4 on FOXO1, SOX4 was knockout and the expression of FOXO1 was overtly decreased accompanied with compromised decidualization as marked by lessened IGFBP1 during decidualization from days 2 to 6 (Fig. 4Μ, Ν). On the other hand, FOXO1 mRNA and protein were upregulated in SOX4 over-expression stromal cell, as well as IGFBP1 protein (Fig. 4O, P). In summary, these results suggested that SOX4 directly regulated transcription of decidualization maker PRL, transcription factor FOXO1, as well as other molecules conducive for human endometrial decidualization.

### SOX4 stabilizes PGR through repressing ubiquitin-proteasome pathway

Previous studies showed that P4-PGR signaling was critical for both initiation and maintenance of decidualization, which incited us to assess PGR expression in the absence of SOX4 during the decidualization. We were attracted by the finding that PGR protein levels were dramatically decreased in the absence of SOX4 on days 0, 2, 3 and 6 during decidualization without overt transcription reduction (Fig. 5A, 5B). Similar results were also observed in primary stromal cells, implying that SOX4 stabled PGR protein at post-transcriptional level (Fig. S3A, S3B). Furthermore, treatment of cells with the proteasome inhibitor MG-132 for 6 hours efficiently attenuated the rapid degradation of PGR protein in SOX4 deficient cells (Fig. 5C). Likewise, PGR protein degeneration was faster in SOX4 depleted hESC two days after EPC treatment in the presence of the protein synthesis inhibitor cycloheximide (CHX) for 0, 3, 6, and 9 h (Fig. 5D). Thus, these results revealed that SOX4 knockdown led to shortened the half-life of PGR protein.

**Fig. 5.**
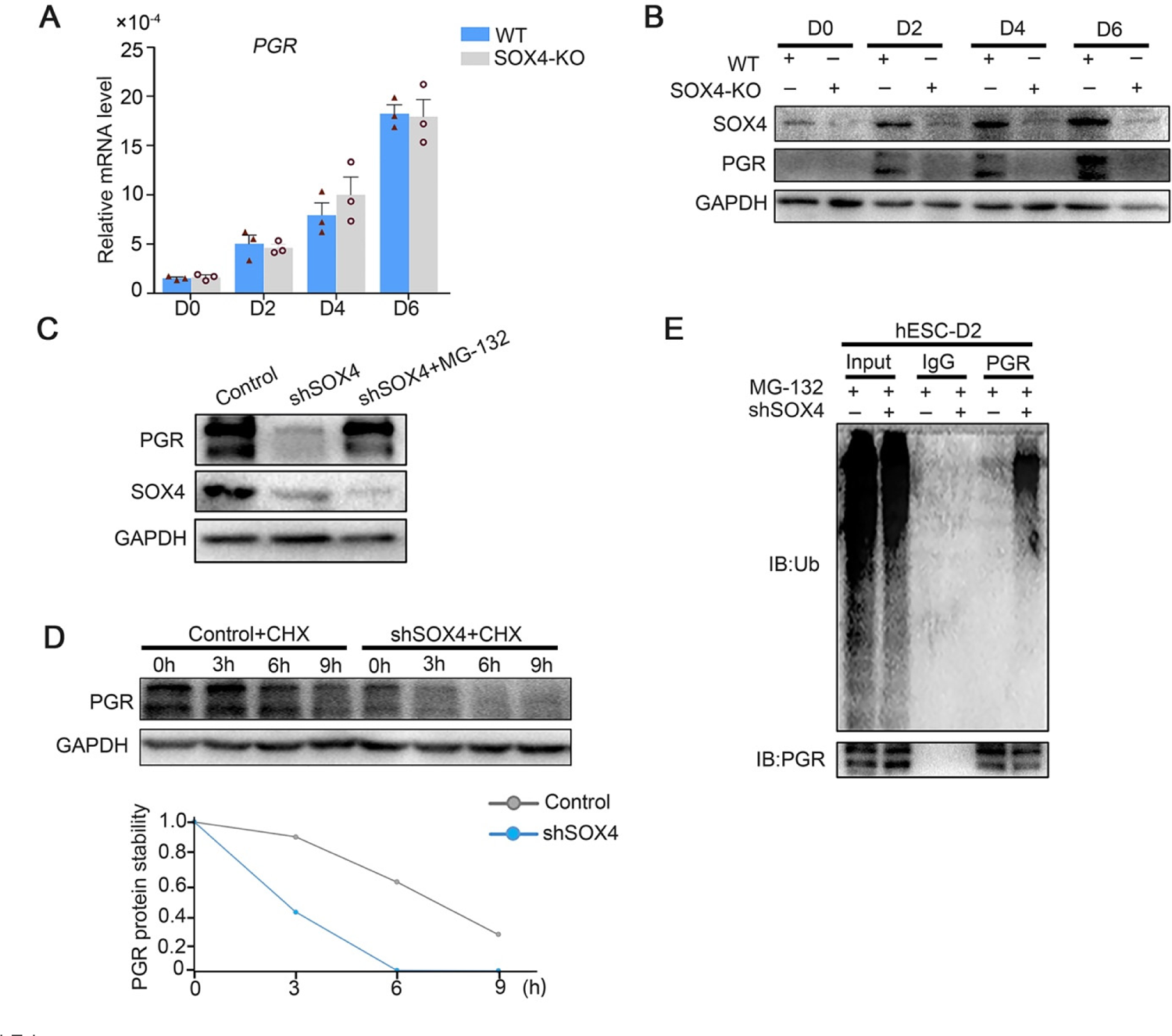
SOX4 stabilizes PGR through repressing the ubiquitin-proteasome pathway. (A) mRNA levels of *PGR* after SOX4 knockout in decidualized immortalized hESCs from days 0 to 6. Results were presented as means ± SEM; n =3. (B) Protein levels of PGR and SOX4 after SOX4 knockout in decidualized hESCs from days 0 to 6. (C) Protein levels of PGR and SOX4 in the presence of shSOX4 or MG132. MG-132 is added 6 hours before collecting the immortalized cells. (D) Protein levels of PGR in the presence of protein synthesis inhibitor cycloheximide (CHX) in SOX4 knockdown cells. (E) PGR ubiquitination after immunoprecipitation with PGR and blotted with ubiquitin antibody in SOX4 knockdown immortalized hESCs decidualized for 2 days. Cells were treated with MG-132 before cell lysis.

To further explore the regulatory apparatus of PGR protein stability, the ubiquitination status of PGR was determined. Immunoprecipitated PGR was blotted against ubiquitin antibody in decidualized immortalized hESCs pretreated with MG-132 for 6 h to postpone protein degradation. Notably, PGR ubiquitination was increased after SOX4 depletion (Fig. 5E). These results suggested that SOX4 knockdown affected PGR protein stability through the polyubiquitin-mediated proteasome degradation pathway.

### SOX4 inhibits PGR degradation by regulating E3 ligase HERC4

Since ubiquitin E3 ligases were mainly responsible for protein ubiquitination, we next intended to identify the potential ubiquitin E3 ligase for PGR protein. Mass spectrometry (MS) was employed for PGR immunoprecipitated (IP) proteins in decidualized immortalized hESCs to estimate PGR associated proteins (Fig. S4A). Reassuringly, several E3 ubiquitination ligases were identified, including HERC4, RNF213 and UBR3 whose expressions were also upregulated according to RNA-Seq in SOX4-silenced hESCs (Fig 6A). Real-time PCR confirmed the abnormal upregulation of these E3 ligases when SOX4 was downregulated (Fig. 6B, and Fig S4B, S4C). The protein level of HERC4 was consistently upregulated upon the ablation of SOX4 (Fig. 6C). As HERC4 showed the most significant change after SOX4 knockdown and its interaction with PGR as a potential E3 ubiquitin ligase was also supported by UbiBrowser database (http://ubibrowser.ncpsb.org), we mainly focused on this E3 ligase in our following experiments.

**Fig. 6.**
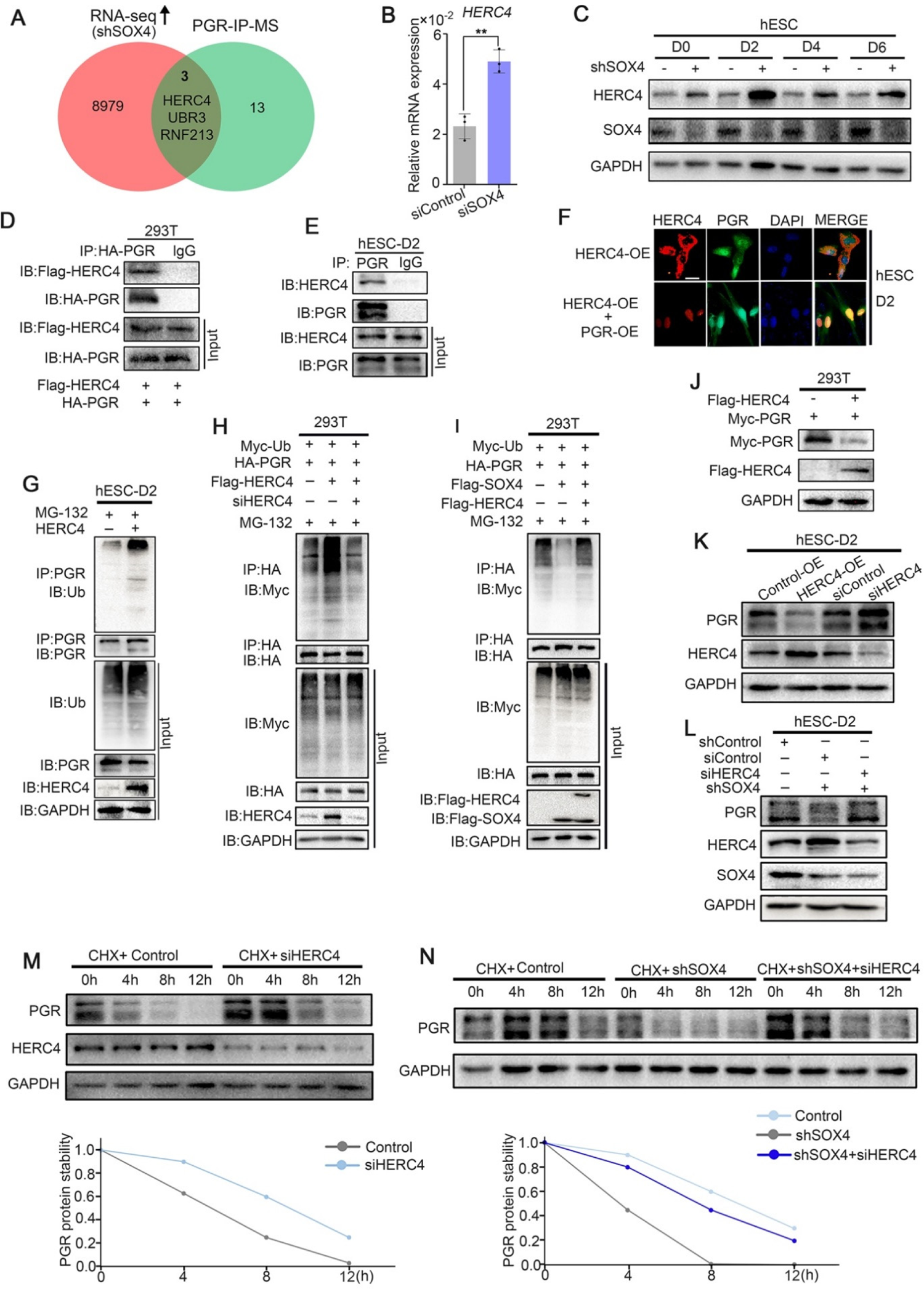
SOX4 inhibits PGR degradation by regulating E3 ligase HERC4. (A) Venn-diagram of overlapping genes between SOX4 up-regulated genes in SOX4 knockdown from RNA-Seq and PGR associated protein form IP-MS. (B and C) mRNA and protein levels of HERC4 in the absence of SOX4 by qPCR (B) and Western-blot (C). Results were presented as means ±SEM; n =3. (D) Protein interaction between PGR and HERC4 after overexpression of HA-PGR and Flag-HERC4 in 293T cells. (E) Protein interaction between PGR and HERC4 after overexpression of HA-PGR and Flag-HERC4 in decidualized immortalized hESCs for two days, D2: two days. (F) Localization of PGR and HERC4 after over-expression of PGR and HERC4 in decidualized hESCs for two days, scale bar: 100 μm. (G) PGR ubiquitination at the present of MG-132 after over-expression of HERC4 in decidualized hESCs for two days. (H) PGR ubiquitination at the present of MG-132 and ubiquitin with HERC4 over-expression or knockdown in 293T cells. (I) PGR ubiquitination at the present of MG-132, ubiquitin and SOX4 with or without HERC4 in 293T cells. (J) Degeneration of PGR after HERC4 overexpression in 293T cells. (K) PGR protein levels at the present of HERC4 overexpression in HERC4 knockdown decidualized immortalized hESCs for 2 days. (L) Protein levels of PGR, HERC4 and SOX4 after SOX4 and/or HERC4 knockdown in decidualized immortalized hESCs for 2 days. (M) The protein half-life of PGR and HERC4 after HERC4 knockdown at the present of protein synthesis inhibitor cycloheximide (CHX) in 2 days decidualized immortalized hESCs. (N) The protein half-life of PGR after SOX4 and/or HERC4 knockdown at the present of protein synthesis inhibitor cycloheximide (CHX) in 2 days decidualized immortalized hESCs. The above experiments were repeated three times.

To verify whether HERC4 was a potential E3 ligase for PGR protein, we first detected the protein interaction between HERC4 and PGR. The co-immunoprecipitation result demonstrated a conserved physical interaction between endogenous HERC4 and PGR in decidualized stromal cells as well as in 293T cells (Fig. 6D, E). Immunofluorescence analysis showed the colocalization of HERC4 and PGR in decidualized hESCs (Fig. 6F). To figure out whether HERC4 could induce endogenous ubiquitination of PGR, HERC4 was overexpressed in immortalized hESCs followed by decidualization for two days. The ubiquitination of PGR was higher in the presence of HERC4 than in control, as shown in Figure 6G, consistent with a reduced level of PGR protein in immortalized hESCs cells (Fig. S4D). Conversely, HERC4 abolished by siRNA significantly reduced PGR ubiquitination in 293T cells (Fig. 6H). The above studies demonstrated that HERC4 was a critical E3 ligase for PGR ubiquitination. Moreover, overexpression of HERC4 recused the decreasing PGR ubiquitination in the presence of SOX4 in 293T cells (Fig. 6I). Hence, SOX4 affected PGR ubiquitination through E3 ubiquitin ligase HERC4.

Since HERC4 was a ubiquitin ligase of PGR and the increased ubiquitination of PGR was observed in the presence of HERC4, we next interrogated whether HERC4 mediated PGR degradation. This posit was underpinned by decreased PGR protein level with HERC4 overexpression in 293T cells (Fig. 6J) and immortalized hESCs (Fig. 6K). PGR protein was also rescued by siRNA-mediated knockdown of HERC4 (Fig. S4E) in SOX4 abolished decidualized immortalized hESCs (Fig. 6L). Likewise, PGR half-life increased in both normal and SOX4 abolished immortalized hESCs after HERC4 repression (Fig. 6M and 6N). In a word, these studies demonstrated that SOX4 mediated PGR degradation by modulating E3 ligase HERC4.

Since there were 41 lysine residues in the PGR protein, UbiBrowser (http://ubibrowser.ncpsb.org) was applied to predict the potential domain recognized by E3 ligase HERC4 (Fig. S5A). As the latent recognition domains were located in DBD and LBD domains of PGR, PGR was divided into four fragments (F1, F2, F3, and F4) accordingly (Fig. 7A). Only F3 (DBD domains of PGR) exhibited protein degradation after co-transfected with HERC4 in 293T cells (Fig. 7B). Point mutation of all Lysine(K) to arginine(R) in DBD region (Fig. 7C) showed that K588, K613, K617, K638 were critical for PGR degradation (Fig. 7D), which was further sustained by incapable PGR ubiquitination mediated by HERC4 in these mutant forms when compared with WT or K565R in 293T cells (Fig. 7E). Collective, these results suggested that K588, K613, K617 and K638 were vital for PGR ubiquitination by HERC4.

### Aberrantly decreased endometrial SOX4 expression is associated with EMS-RIF undergoing IVF treatment

There was considerable evidence indicating that hESCs decidualization was severely impaired in patients with the EMS, both in eutopic and ectopic lesions(Klemmt, Carver et al. 2006). RIF has also been reported to be associated with compromised decidualization. We next analyzed the expression levels of SOX4, PGR, FOXO1, HERC4, IGFBP1, and PRL in the mid-secretory endometrium of normal women (control, n = 12) versus women who suffered RIF due to endometriosis (EMS-RIF, n = 12). The expressions of *SOX4*, *FOXO1*, *IGFBP1*, and *PRL* were significantly decreased in EMS-RIF group with increased HERC4 compared with the control, but the mRNA level of *PGR* was comparable in both groups (Fig. 8A–F). At protein level, a large portion of endometrial samples from women with EMS-RIF showed reduced expression of SOX4, FOXO1, IGFBP1. Although the *PGR* mRNA levels were comparable between normal and RIF samples, PGR protein was significantly decreased accompanied by increased HERC4 in EMS-RIF group (Fig. 8G, H). Immunostaining analysis further revealed significantly reduced PGR, SOX4, IGFBP1 and FOXO1 expression and increased HERC4 in stromal cells in women with EMS-RIF (Fig. 8I).

**Fig.7.**
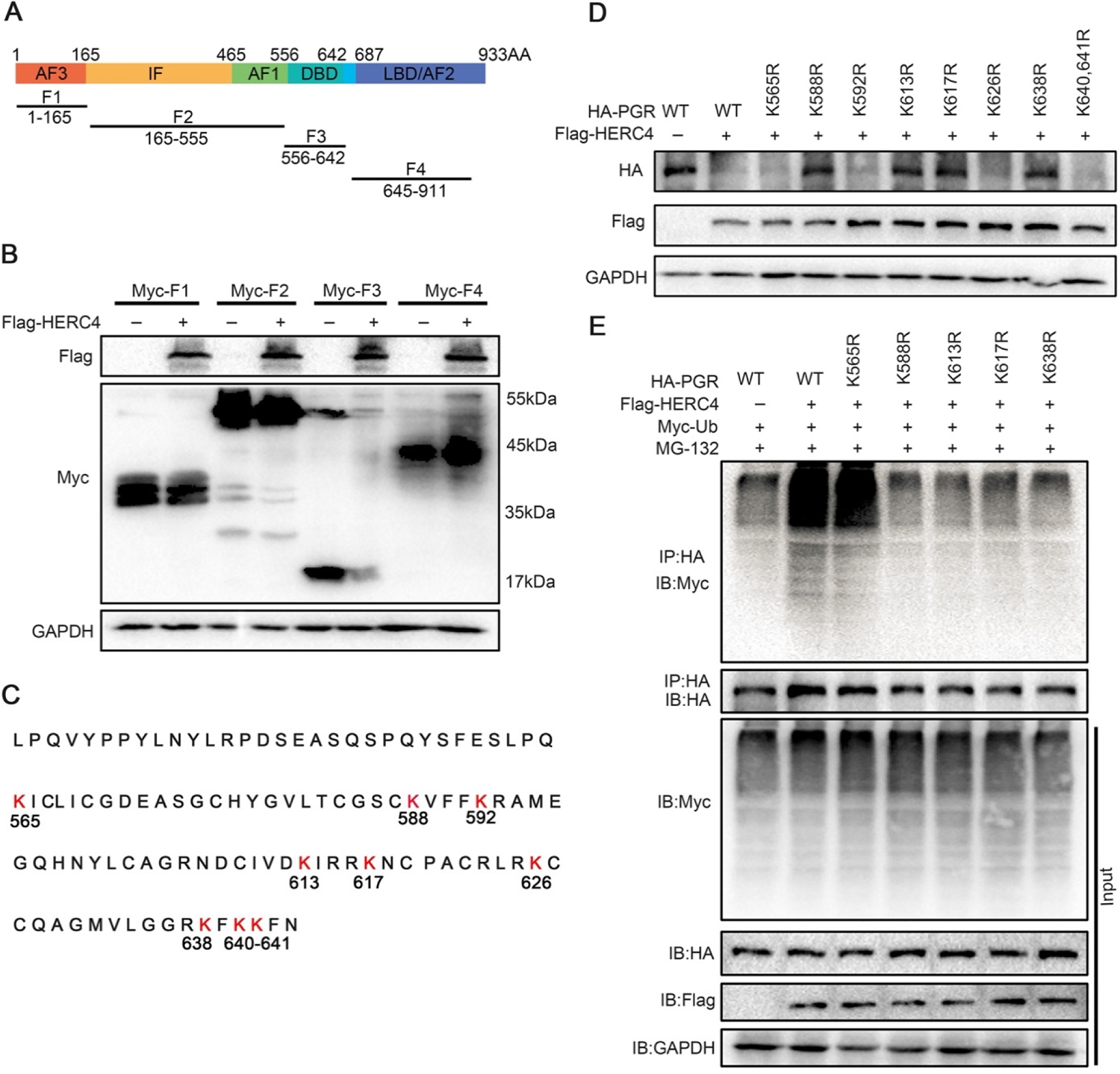
HERC4 mediates PGR ubiquitination at K588, K613, K617 and K638. (A) Schematic diagram of PGR structure. F1 to F4 represents four different functional domains, respectively. (B) Protein levels of different fragments of PGR (Myc tagged F1-F4) at the present of HERC4 in 293T cells. (C) Lysine (K) sites in PGR DBD domain. (D) Protein levels of PGR after Lysine mutant to Arginine in DBD with HERC4 over-expression in 293T cells. (E) PGR ubiquitination after Lysine mutant to Arginine in DBD at the present of HERC4, ubiquitin and MG-132 in 293T cells.

**Fig. 8.**
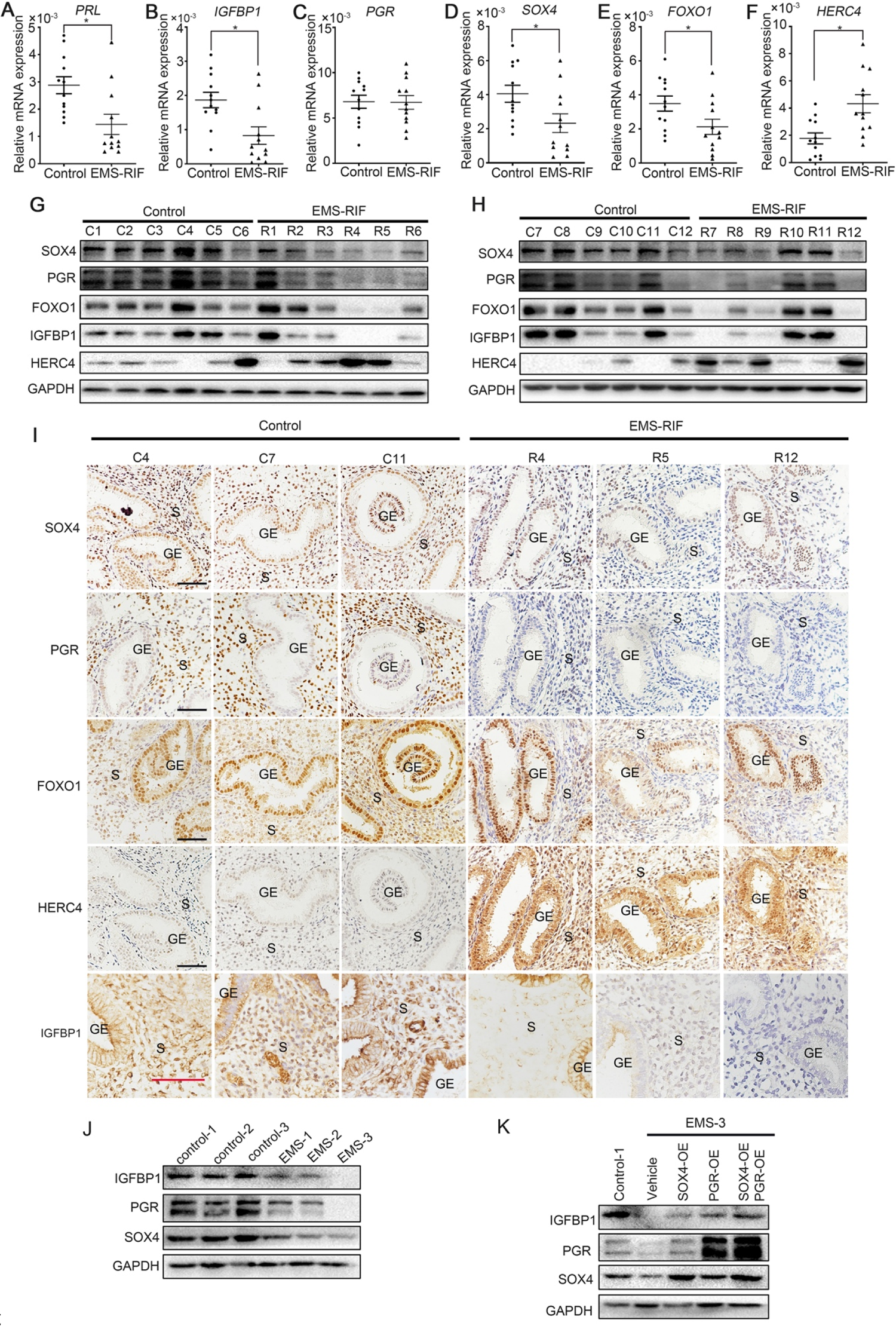
Endometrial SOX4 expression in RIF of women with EMS undergoing IVF treatment. (A, B, C, D, E, and F) Expression of *SOX4*, *FOXO1*, *IGFBP1*, *PRL*, *PGR*, and *HERC4* in mid-secretory endometrium from control (n=12) and EMS-RIF (n = 12). Results were presented as means ±SEM. * indicates P<0.05. (G and H) Protein levels of SOX4, FOXO1, IGFBP1, PGR, and HERC4 in mid-secretory endometrium from control (n=12) and EMS-RIF (n = 12). C1-C12 and R1-R12 represent tissues from different patient. (I) Localization of SOX4, FOXO1, IGFBP1, PGR, and HERC4 in control (n = 3) and EMS-RIF groups as detected by immunostaining. C4, C7, C11, R4, R5 and R12 represent tissues from different patient. GE: Gland epithelium; S: Stromal. Scale bar: 100 μm. (J) Protein levels of SOX4, IGFBP1, and PGR in primary hESCs of control and EMS by Western-blot. (K) Protein levels of IGFBP1, PGR and SOX4 in decidualized primary endometrial stromal cells from EMS-3 after overexpression of SOX4 and/or PGR.

Additionally, primary human endometrial stromal cells (hESCs) were obtained from the proliferative phase endometrium of three normal and three EMS-RIF patients. The expression levels SOX4, PGR, IGFBP1 were lower in the primary hESCs of EMS-RIF cultured with EPC for 4 days compared to the control (Fig. 8J). Moreover, overexpression of SOX4 and/or PGR restored IGFBP1 expression in EMS-RIF stromal cells (Fig. 8K). Collectively, our in vivo and in vitro evidence strongly suggested the crucial role of SOX4 in decidualization and female fertility.

## Discussion

Here, we find that SOX4 is the most abundant SOX family member in hESCs and plays a vital role in decidualization. Previous studies have demonstrated that the SOX family controls cell fate and differentiation in various developmental processes(She and Yang 2015, Angelozzi and Lefebvre 2019). A previous study has confirmed that epithelial SOX17, one of SOX transcription factors, is indispensable for embryo implantation by regulating IHH expression(Wang, Li et al. 2018). SOX17 is also a crucial transcription factor located in the endothelium, especially in artery, regulating permeability of blood vessel(Corada, Orsenigo et al. 2013). Since SOX17 also abundantly expresses in uterine endothelial cells, the physiological role of endothelial SOX17 during pregnancy remains to be explored.

In the present study, we uncover the critical role of SOX4 in stromal cell decidualization by RNA-Seq and ChIP-Seq. Several previously unappreciated genes of SOX4 are unveiled through RNA-seq. FOXO1 is indispensable in the process of decidualization account for the widely overlapping of binding peaks with PGR(Vasquez, Mazur et al. 2015). In this investigation, FOXO1 and PRL are regulated by SOX4 at transcriptional level, indicating the essential role of SOX4 in decidualization. We are also very surprised to notice that the direct regulation of SOX4 on FOSL2, an important part of AP1 complex involving inflammation, expression directly. Moreover, we also find that FOSL2 and FRA2 motif are significantly enriched in SOX4 binding sites, suggesting the complex interplay between these factors. The regulation of SOX4 on FOLS2 implicates an alternative role of SOX4 regulating decidualization, which deserves further investigation.

PGR is a master regulator of the decidualization process since decidualization is mainly influenced by the progesterone-PGR signaling. After dimerization and nuclear translocation, PGR protein interacts with the following proteins to synergistically direct the differentiation program: FOXO1, C/EBPβ, STAT3, STAT5, HOXO10, and HOXO11(Gellersen and Brosens 2014). An array of modulators regulates the functional plasticity of PGR, including subcellular distribution, protein modification, and interaction with co-regulators(Wu, Li et al. 2018). In this study, we were intrigued by the finding that SOX4 affects PGR protein stability rather than its transcription.

During the menstrual cycle, estrogen-ER signaling induced endometrial PGR mRNA in the proliferative phase, recapitulating the regulation manner of PGR in mouse uteri(Tan, Paria et al. 1999, McKinnon, Mueller et al. 2018). The PGR protein is further stabilized by increased SOX4 at post-transcriptional regulation in the later secretory phase. Previous studies have validated that SOX4 regulates P53 stability through E3 ligase Mdm2(Pan, Zhao et al. 2009). In breast cancer cells, E3 ligase BRCA1(Calvo and Beato 2011) and CUEDC2(Zhang, Zhao et al. 2007) have been shown to regulate PGR protein stability. Multiple evidence in our research have verified that the E3 ligase HERC4 functions as a ubiquitin-modified enzyme for the PGR protein. Further exploration revealed that lysine residual in the PGR DNA-binding domain possess modified sites for ubiquitination. In the endometrial cells, there are very limited reports regarding PGR protein modification. Our previous study proved that Bmi1 facilitates the PGR ubiquitination through E3 ligase E6AP, which promotes PGR transcriptional activity instead of protein degradation(Xin, Kong et al. 2018). Besides, SUMOlation of PGR has been reported to occur at the K388 site and fine-tunes the transcriptional activity of PGR(Jones, Fusi et al. 2006). The Lys-388 is a key site not only for PGR-B SUMOlation, but also a key site for CUEDC2-mediated ubiquitination in proteasome in breast cancer. Here, we characterized that K588, K613, K617, and K638 were critical sites for E3 ligase HERC4, which mediated PGR ubiquitination and degradation. There were other E3 ligases interacting with PGR as revealed by our mass spectrum, whether these E3 ligases also mediated PGR ubiquitination remains largely unknown. Multiple specific E3 ligases may exist for PGR ubiquitination rely on individual lysine residues.

Here, we report a previously unrecognized PGR ubiquitination and degradation modified by the ubiquitin E3 ligase HERC4 regulated by SOX4 in endometrial stromal cells. There are several potential postulations concerning the underlying mechanism by which SOX4 inhibits HERC4 expression in stromal cells. SOX4 has been reported to interact with repressive histone modifiers, such as H3K27 trimethylation enzyme EZH2, to directly repress the target gene transcription(Koumangoye, Andl et al. 2015). On the other hand, SOX4 indirectly repress the target gene expression through regulating EZH2 expression(Tiwari, Tiwari et al. 2013). Since no direct SOX4 binding on HERC4 is observed, the detailed mechanism by which SOX4 regulates HERC4 transcription in endometrial stroma cells requires further exploration.

Considering the critical role of progesterone during the pregnancy establishment and maintenances, both natural and synthetic progestogens have been widely used to improve endometrial function in women with a history of RIF and unexplained recurrent pregnancy loss. However, some of these patients still suffered from RIF and recurrent miscarriages due to progesterone resistance. The causes of progesterone resistance may be related to defects in the PGR signaling and its molecular chaperones as well as decreasing PGR transcriptional activity(McKinnon, Mueller et al. 2018). PGR functional deficiency is related to abnormal PGR mRNA levels, post-transcriptional modifications, post-translational modifications and protein stability(Xin, Qiu et al. 2016, McKinnon, Mueller et al. 2018). However, the specific causes of PGR defects are, at present, poorly explored and understood.

Endometriosis is frequently accompanied by progesterone resistance and infertility. It is interesting that lower SOX4 expression is strongly relevant with reduced PGR protein expression in endometrial stromal cells of EMS who experience RIF, and the level of PGR is restored to some extent when SOX4 is overexpressed in these stromal cells. It is conceivable that PGR defects caused by insufficient SOX4 may potentially result in implantation failures, even with high doses of progestogens supplementation during IVF. In our previous study using human endometrial samples from different patients with recurrent spontaneous abortion cohort, we also observed progesterone resistance as revealed by defective PGR-signaling, with normal PGR protein level but diminished transcriptional PGR activity(Xin, Kong et al. 2018). This implies that any disturbances in the transduction of P4-PGR signaling pathway will severely influence the endometrial cell responsiveness to progesterone and contribute to infertility.

Collectively, our investigation provides compelling evidence that SOX4 plays a key role in hESCs decidualization through the directly transcriptionally regulation of decidual maker PRL and many other critical factors related to decidualization. Meanwhile, SOX4 endows human stromal cells appropriate progesterone responsiveness by fine-tuning PGR protein stability. Aberrant SOX4 expression is strongly associated with decreased PGR and FOXO1 expression in the endometrium of women who have experienced RIF with EMS, implying the high clinic relevance of SOX4 in female fertility and pregnancy maintenance. A better understanding of the regulatory network of SOX4 will facilitate the development of therapeutic strategy for the clinical treatment of RIF in EMS.

## Materials and Method

### Sample of clinical cases

Healthy and infertile women with RIF due to endometriosis were recruited from Liuzhou Maternity and Child Health Care Hospital in China from March 2018 to December 2020. The study had been approved by hospital ethics committee and all participants signed informed consent. The age of participates are between 20 and 38 years of age with Body Mass Index (BMI) between 18 and 23. The thickness of the endometrium on ovulation days was between 8 and 16 mm with menstrual cycle between 28 ± 7 days and no steroid treatment or other medication for at least 2–3 months before biopsy. Detailed information of patients was listed in Supplementary Table 2. Patients with polycystic ovary syndrome, endometrial polyps, chronic endometritis, and hydrosalpinges were excluded. Only endometrial tissue with no apparent pathology assessed by a pathologist was kept for further experiments. Recurrent implantation failure (RIF) referred to the inability to achieve a clinical pregnancy after transferring at least four good-quality embryos in a minimum of three fresh or frozen cycles in a woman under 40 years old(Coughlan, Ledger et al. 2014).

### Isolation and culturing of primary endometrial stromal cells

Primary hESCs were obtained from healthy, reproductive-aged volunteers with regular menstrual cycles or endometriotic patients with infertility. An endometrial biopsy was performed during the proliferative phase of the menstrual cycle. These participants were recruited from The First Affiliated Hospital of Xiamen University in China from July 2019 to October 2020. The study had been approved by hospital ethics committee and all participants signed informed consent. Participants were documented not under hormone treatments for at least 3 months before surgery. Only endometrial tissue with no apparent pathology assessed by a pathologist was kept for further experiments. Primary human endometrial stromal cells (hESCs) of endometriosis women were isolated by laparoscopy, and the isolation and culturing of primary endometrial stromal cells were performed as follows. The endometrial tissues were first cut into pieces as small as possible and subjected to type IV collagenase (2% concentration) digestion for 1h. Two hours after cell seeding, culture medium was changed to remove the floating cells.

### Establishment of SOX4 knockout cell lines

Immortalized hESC cell lines were purchased from ATCC Corporation (American Type Culture Collection, ATCC® crl-4003^TM^). SOX4 knockout was generated by CRISPR/Cas9 approach as previously described(Zhang, Meng et al. 2019). (1) sgRNAs targeting SOX4 gene (NM_003107.3) were designed on the website (https://zlab.bio/guide-design-resources) and subcloned into the pL-CRISPR.EFS.GFP vector (Addgene plasmid #57818). The target sequences were listed in Supplementary Table 1. (2) Cas9 plasmid was packaged with lentivirus. After infecting stromal cells, Cas9 positive cells were sorted and were plated into a 96-well plate with one cell per well. (3) knockout efficiency was verified by DNA sequencing and Western blot after single cell derived clone expansion.

### In vitro culture of hESCs and decidualization

Immortalized hESCs were maintained in DMEM/F12 without phenolic red (Gibco) in the presence of 10%charcoal stripped fetal bovine serum (CS-FBS, Biological Industries), glucose (3.1 g/L), sodium pyruvate (1 mM, Sigma); sodium bicarbonate (1.5 g/L, Sigma), Penicillin-Streptomycin (50 mg/ml, Solarbio); insulin-transferrin-selenium (ITS, 1%, Thermo Fisher) and puromycin (500 ng/ml). The primary cultured cells were maintained in DMEM/F12 (Gibco) without phenolic red and 10% CS-FBS. For decidualization, hESCs were cultured in medium at the present of estrogen (E2, 10nM, Sigma), medroxyprogesterone acetate (MPA, 1 mM, Sigma), and dibutyl cyclophospsinoside (db-cAMP, 0.5 mM, MCE) in 2% CS-FBS with different days. All the cells were cultured in 5% CO_2_, 95% air, 100% humidity at 37 ℃ and culture medium was replaced every 48 h.

### Plasmid construction and siRNA transfection

The overexpressed plasmids of SOX4 (pLVML-FLAG-SOX4-IRES-puro, and pLVX-HA-IRES-ZSgrenn-SOX4), HERC4 (pLVX-FLAG-HERC4-IRES-Zsgreen, pENTER-FLAG-HERC4, and HA-PCMV-HERC4), PGR (pLVX-MYC-PGR-IRES-Zsgreen and HA-pCMV-PGR), and ubiquitin (pRK5-HA-Ubiquitin-WT and MYC-Ubiquitin-WT) were purchased from Wu Han Miao Ling Company or home-made. hESCs were transfected with these plasmids by Lipofectamine 2000 transfection reagent (Invitrogen) or infected with the lentivirus. ShSOX4 lentivirus was purchased from Shanghai Ji Man Company. The ShSOX4 lentivirus and the control lentivirus were placed in the hESC medium, 48h after infection, hESCs were decidualized with EPC. siRNAs targeting SOX4, HERC4, PGR and FOXO1 were purchased from Guangzhou RiboBio Biological Company (see the specific sequence for details in Supplementary Table 1). RNA interference was carried out according to the manufacturer’s instructions. Briefly, 10 mM siRNA was transfected into hESC with Lipofectamine RNAi MAX (Invitrogen, Carlsbad, USA).

### ChIP-seq, ChIP-QPCR

ChIP-seq was performed according to manual of ChIP-IT high Sensitivity (Active motif, catalog NO. 53040). Briefly immortalized hESCs (treated with EPC for two days) with exogenous expressed HA-SOX4 were cross-linked and immunoprecipitated with HA (CST, c29F4) and PGR (CST, 8757) antibodies. Immunoprecipitated and input DNA were quantified using Qubit 4.0 fluorometer. Libraries were prepared using the KAPA DNA HyperPrep Kit (KK8502) and sequenced with an Illumina Nova-PE150. After filter the raw data to remove adapter from read by Trimgalore, the clean reads were aligned to the human genome (Hg38) by HISAT2. Only uniquely aligned reads were kept for downstream analysis. MACS2 was applied for peak call using default parameters. The enrichment of SOX4 binding on specific gene was visualized in Integrative Genomics Viewer (IGV). All the PCR primers used in ChIP-qPCR were listed in Supplementary Table 1.

### Immunoprecipitation

For protein interaction, IP experiments were performed as previously described(Xin, Kong et al. 2018). Anti-PGR (CST, 8757), Anti-HA (CST, c29F4), Anti-Myc (CST, 2276), Anti-Flag (Sigma, F1804) and mouse IgG or rabbit IgG (Mouse Anti-Rabbit IgG (Conformation Specific, L27A9 mAb) were used for immunoprecipitation. The immunoprecipitants were washed four times in lysis buffer, resolved with SDS-PAGE, and immunoblotted with corresponding antibodies.

### RNA-Seq and bioinformatic analysis

Scramble or siSOX4 RNA were transfected into hESCs at 50% confluence with RNAi MAX (Invitrogen). Immortalized hESCs were decidualized for 2 days after transfection and RNAs were collected for RNA-Seq of BGISEQ (China, BGI). Purified RNA was prepared and subjected to 50-bp single-end RNA sequencing. RNA-seq raw data were initially filtered to obtain clean data after quality control by Trimgalore. Clean data were aligned to the human genome (Hg38) by HISAT2. RPKM value of each gene was calculated by EdgeR package in R.

### IP mass spectrometry

Immortalized hESCs were used to prepare IP samples after treated with EPC 2 days. After Co-IP with PGR-conjugated agarose beads, the immunoprecipitants were resolved with SDS-PAGE and visualized using Coomassie brilliant blue stain. The discrete bands between PGR and IgG were isolated, digested, purified, and subjected to Liquid Chromatograph Mass Spectrometer (LC-MS) in School of Life Sciences, Xiamen University.

### Construction of dual-luciferase reporter

Construction of luciferase reporter was performed as previously described(Jiang, Liao et al. 2015). The promoter regions of SOX4 were amplified from genomic DNA and subcloned into pGL3 plasmid. All constructs were transiently transfected into 293T cells using Lipofectamine 2000. Total cell lysates were prepared 36 h after transfection, and luciferase activity was measured using the Dual-Luciferase Reporter Assay System (Promega Corporation). Firefly luciferase activity was normalized by Renilla luciferase activity.

### PGR ubiquitination assay

The ubiquitination assays were performed as previously described(Xin, Kong et al. 2018). MG-132 (Sigma M8699, 20 μM) was added to the cultured medium 6 h before cells collection. Cell lysis was immunoprecipitated with antibodies against PGR, HA and Myc, respectively. Proteins were released from the beads by boiling in SDS-PAGE sample buffer, and ubiquitination was analyzed by immunoblotting with different antibodies.

### Immunostaining

After deparaffinization and hydration, formalin-fixed paraffin embedded endometrial sections (5 μm) were subjected to antigen retrieval by autoclaving in 10 mM sodium citrate solution (pH=6.0) for 10 min. A diaminobenzidine (DAB, Sigma) solution was used to visualize antigens. Sections were counter-stained with hematoxylin. In immunofluorescence studies, formalin-fixed hESC cells were blocked with 5% BSA in PBS and immune-stained by antibodies for SOX4 (Abcam, ab80261), PGR (CST, 8757), FOXO1 (Abcam, ab39670), IGFBP1(Abcam, ab228741) and HERC4(Proteintech,13691-1-AP). Signals were visualized by secondary antibody conjugated with Cy2 or Cy3 fluorophore (Jackson Immunoresearch). Sections were counter-stained with Hoechst 33342 (2 μg/ml, Life technology).

### Western-blot and Real-time PCR

Western blot analysis was performed as described previously(Zhang, Meng et al. 2019). Antibodies against SOX4, IGFBP1, FOXO1, PGR, HERC4, HA, Myc, Flag, and ubiquitin were used. GAPDH served as a loading control. Quantitative real-time PCR was performed as described previously(Jiang, Liao et al. 2015). Total RNA was extracted from hESCs or endometrium using TRIzol reagent (Invitrogen) following the manufacturer’s protocol. A total of 1 μg RNA was used to synthesize cDNA. Quantitative real-time PCR was performed with SYBR Green (Takara) on an ABIQ5 system. All expression values were normalized against GAPDH. All PCR primers are listed in Supplementary Table 1.

## Statistics

Statistical analysis was performed with Graphpad Prism. The data were shown as the mean ± SEM. Statistical analyses were performed using 2-tailed Student’s *t-*test or ANOVA. All experiment were repeat at least three times and *p*-values less than 0.05 were considered statistically significant.

## Data availability

The datasets generated and analyzed in the study are available in the NCBI Gene Expression Omnibus (GEO). RNA-seq and ChIP-Seq data sets generated in this study have been deposited at the Gene Expression Omnibus database under accession number GSE146280 and GSE174602 (token: gdczyosqzzozbcx), respectively. RNA-seq for mouse uterus is from a previous study (GSE116096).

## Author contributions

P.H., W.D., M.L., X.Z. and H.C. performed experiments and prepared figures. P.H., W.D., J.L., H.W., A.Q. and S.K. designed experiments. Z.L., J.W., M.Q., and Y.Y. collected the samples and clinical information. P.H., W.D., and S.K. analyzed data. P.H., W.D., H.B., F.R., J.C., D.C., H.W., A.Q. and S.K. wrote the manuscript. All authors read and approved the final manuscript.

## Acknowledgements

This work was supported by National Key R&D program of China (2017YFC1001402 to H.W., 2018YFC1004404 to S.K.), National Natural Science Foundation of China (81830045 and 82030040 to H.W., 81960280 to A.Q., 82001553 to P.H., 81701483 and 81971419 to W.D.), Fundamental Research Funds for the Central Universities (20720190073 to W.D.) and Guangxi natural science foundation project (2019JJB140179 to P.H.).

## Supplementary Materials

**Supplementary Figure 1.**
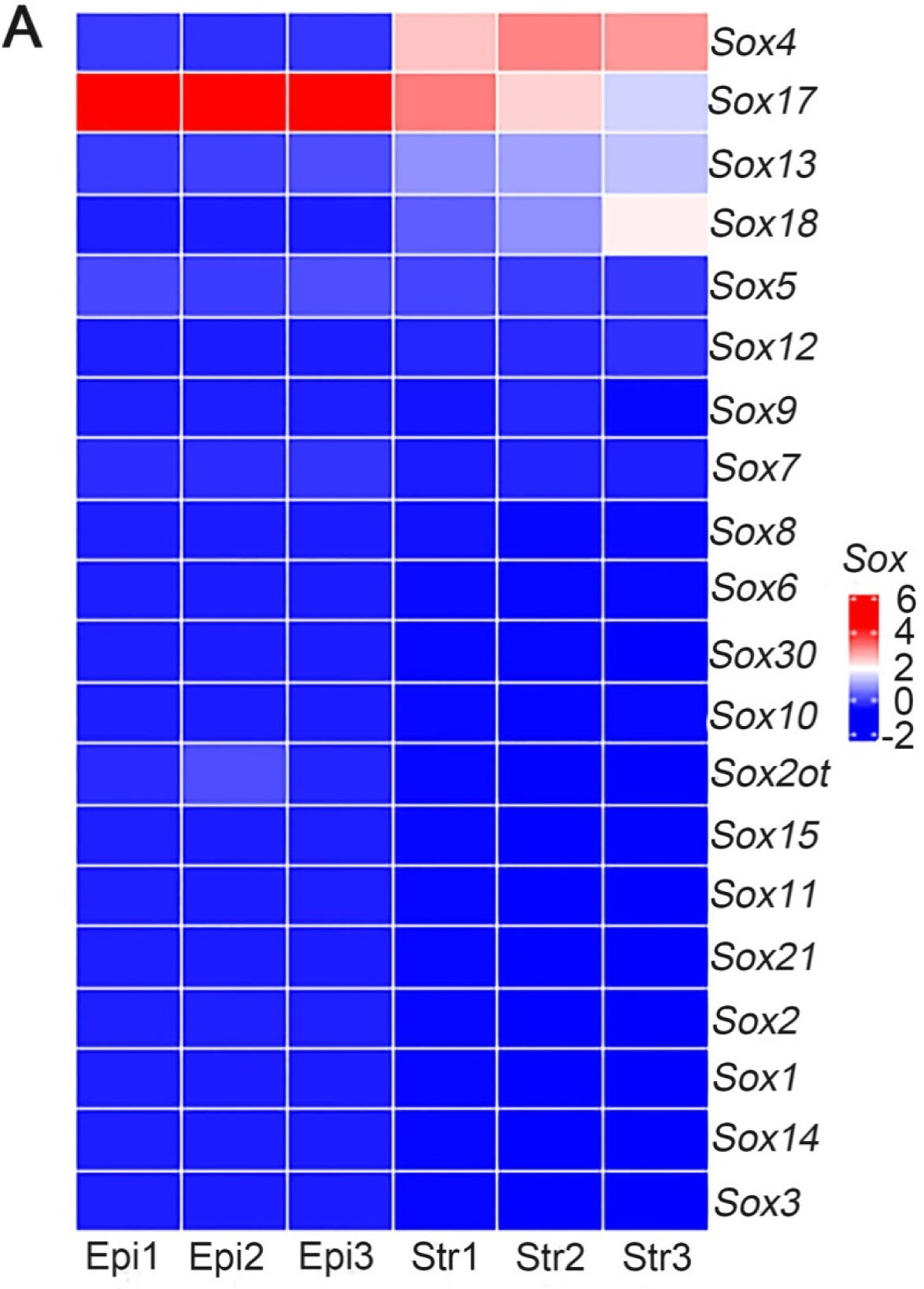
Sox family expression in mouse uterine stromal and epithelial cells. (A) Heatmap of Sox family members in mouse uterine epithelial and stromal cells by RNA-Seq, n = 3.

**Supplementary Figure 2.**
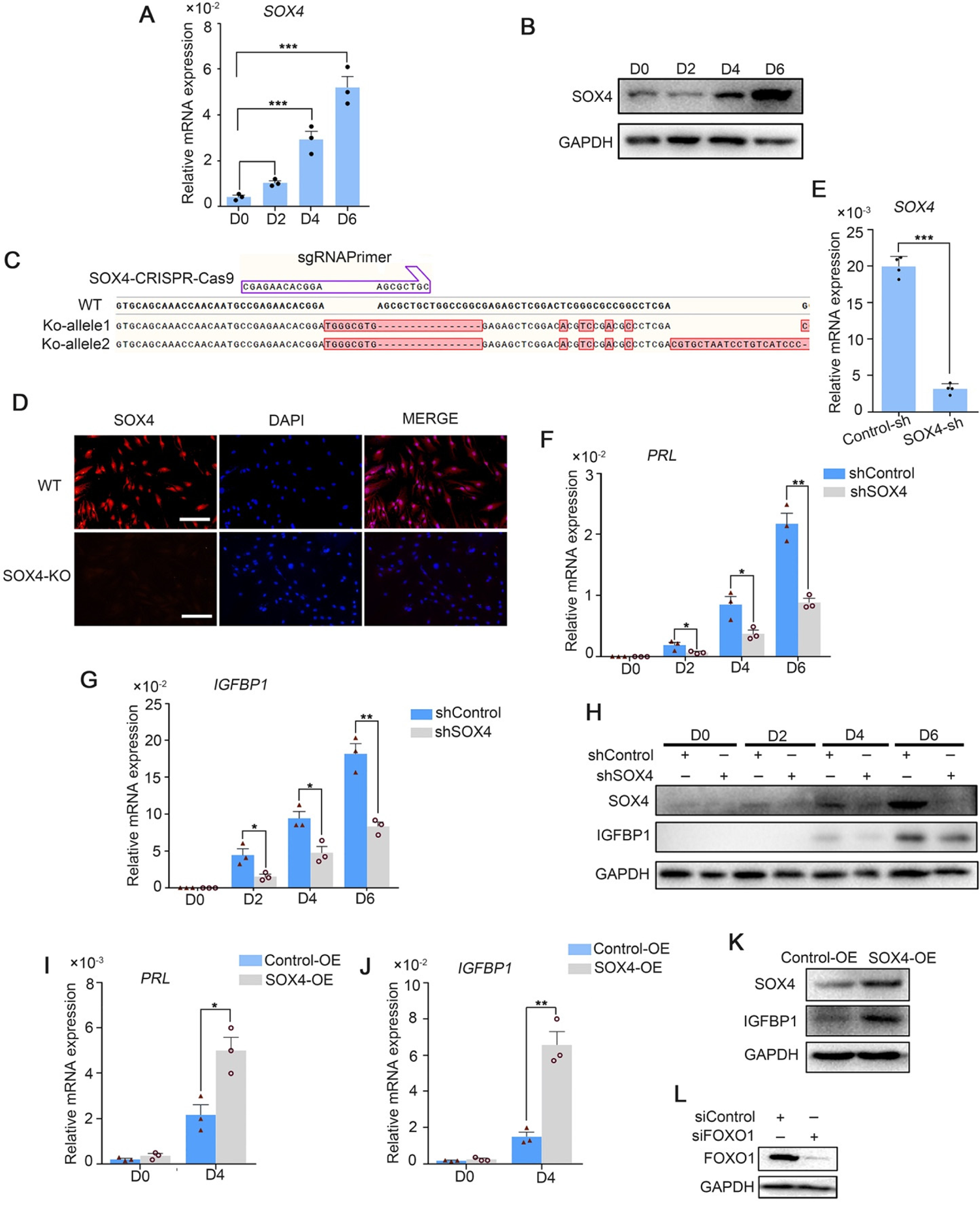
SOX4 is essential for decidualization in primary endometrial stroma cells. (A and B) Expression of SOX4 mRNA and protein in decidualized primary hESCs from day 0 to day 6. Results were presented as means ±SEM; n =3; *** indicates P<0.001. (C) Schematic diagram of SOX4 knockout by CRISPR/Cas9. (D) SOX4 knockout confirmation by immunofluorescence in hESCs. Scale bar: 50 μm. (E) Knockdown efficiency of SOX4 by shRNA. Results were presented as means ±SEM; n =3; *** indicates P<0.001. (F and G) Expression of of *PRL* and *IGFBP1* in the SOX4-knockdown primary decidualized hESCs from days 0 to 6. Results were presented as means ±SEM. n =3; * indicates P<0.05, ** indicates P<0.005. (H) Protein levels of SOX4 and IGFBP1 in the SOX4-knockdown primary decidualized hESCs from days 0 to 6. (I and J) Expression of *PRL* and *IGFBP1* in primary decidualized hESCs for 4 days after SOX4 over-expression. Results were presented as means ±SEM; n =3; * indicates P<0.05, ** indicates P<0.005. (K) SOX4 expression after SOX4 overexpression in the primary decidualized hESCs for four days. (L) FOXO1 expression after FOXO1 knockdown in the primary decidualized hESCs.

**Supplementary Figure 3.**
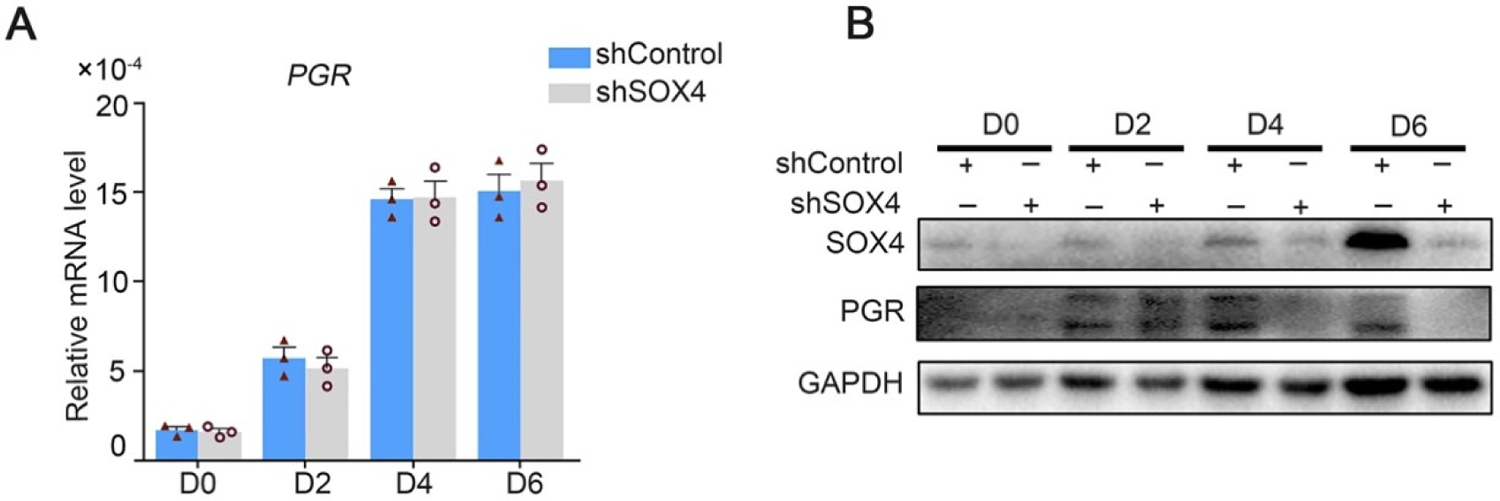
Regulation of SOX4 on PGR in primary decidualized stroma cells. (A) Expression of *PGR* mRNA after SOX4 knockdown in primary decidualized hESCs cultured with EPC from 0–6 days. Results were presented as means ±SEM; n =3. (B) Protein levels of PGR after SOX4 knockdown in primary decidualized hESCs cultured with EPC from 0–6 days.

**Supplementary Figure 4.**
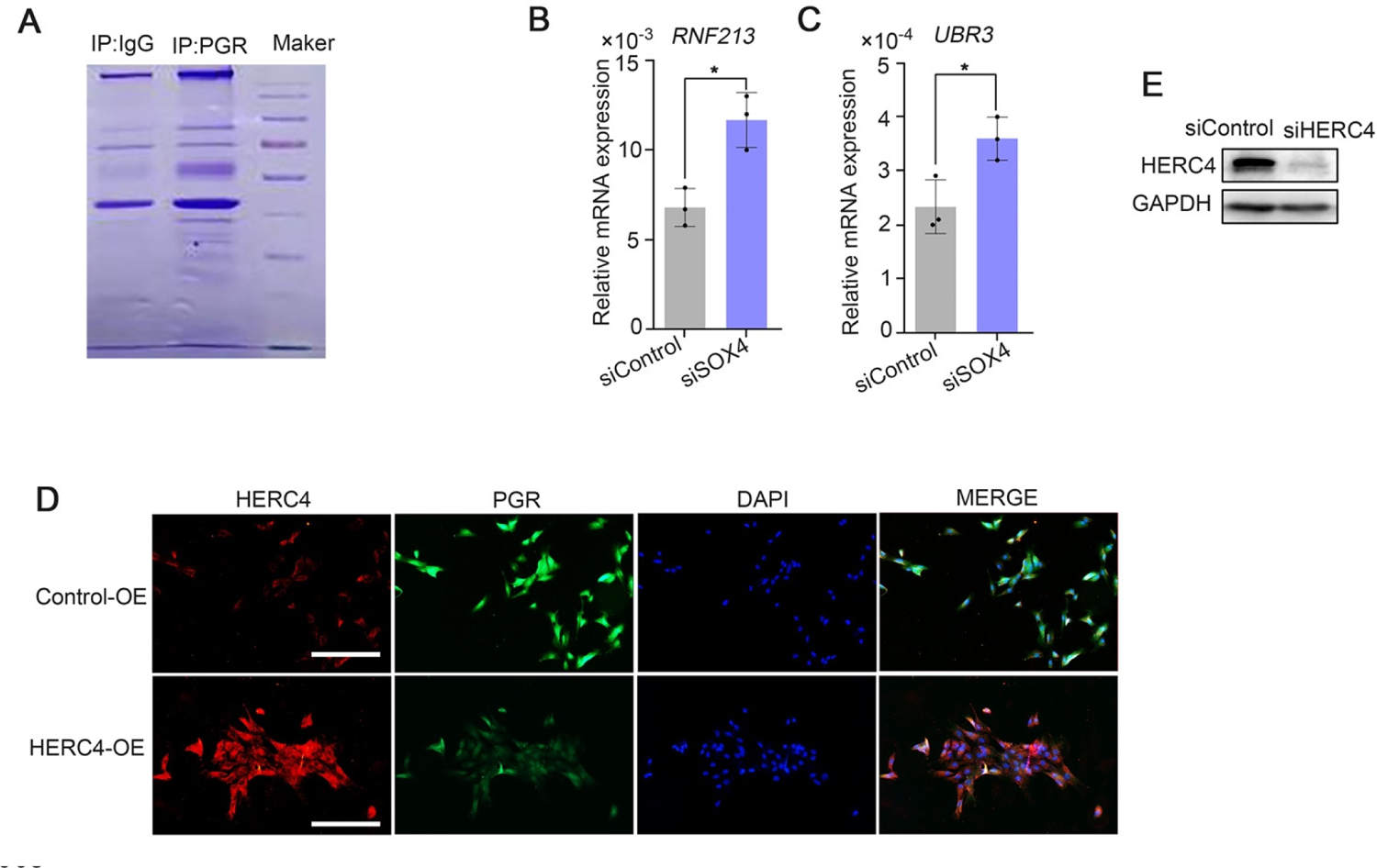
Expression of SOX4 regulated ubiquitin E3 ligase in decidualized hESCs. (A) Coomassie blue staining of PAGE-Gel with PGR immunoprecipitated complex. (B and C) Expression of *RNF213* and *UBR3* in SOX4 knockdown hESCs. Results were presented as means ±SEM; n =3; * indicates P<0.05. (D) PGR expression after HERC4 over-expression in hESCs as detected by immunofluorescence, scale bar: 100 μm. (E) Knockdown efficiency of HERC4 by siHERC4. The above experiments were repeated three times.

**Supplementary Figure 5.**
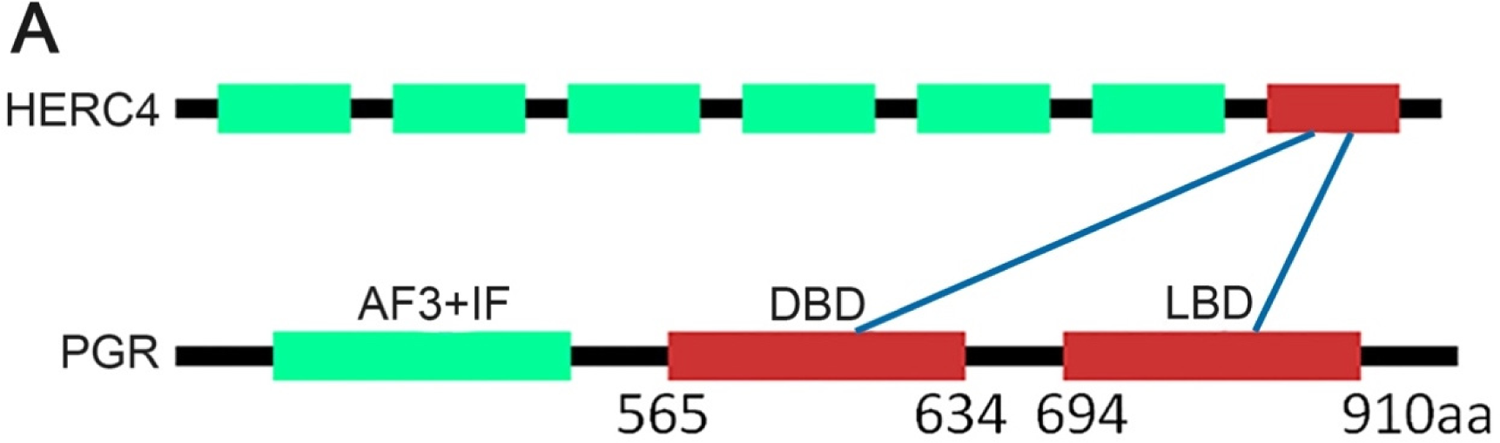
The predicted lysine sites for the ubiquitin modification in PGR protein by the E3 ligase HERC4. (A) UbiBrowser (http://ubibrowser.ncpsb.org) was applied to predict the potential site recognized by E3 ligase HERC4.

**Supplementary Table 1:**
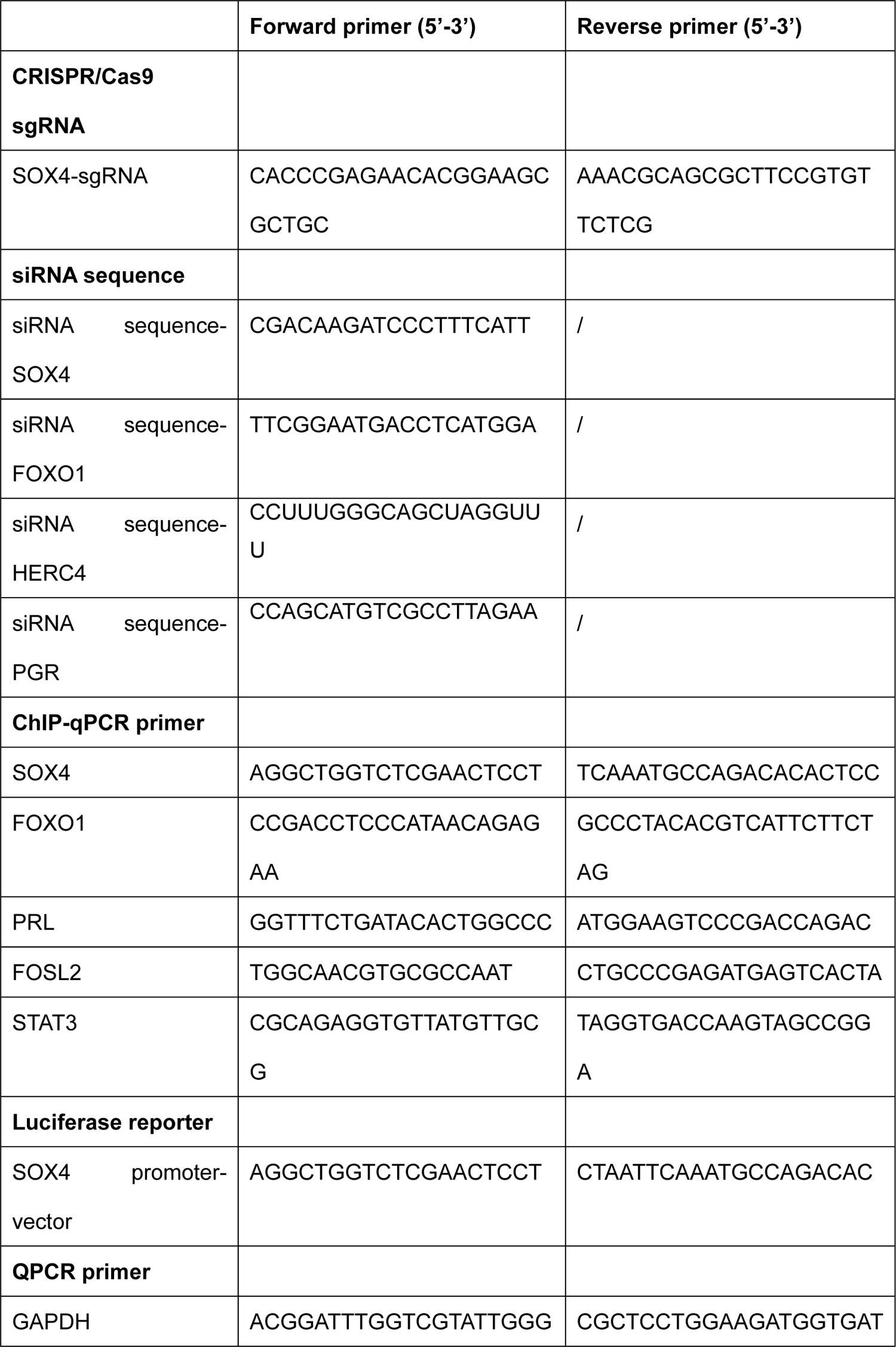
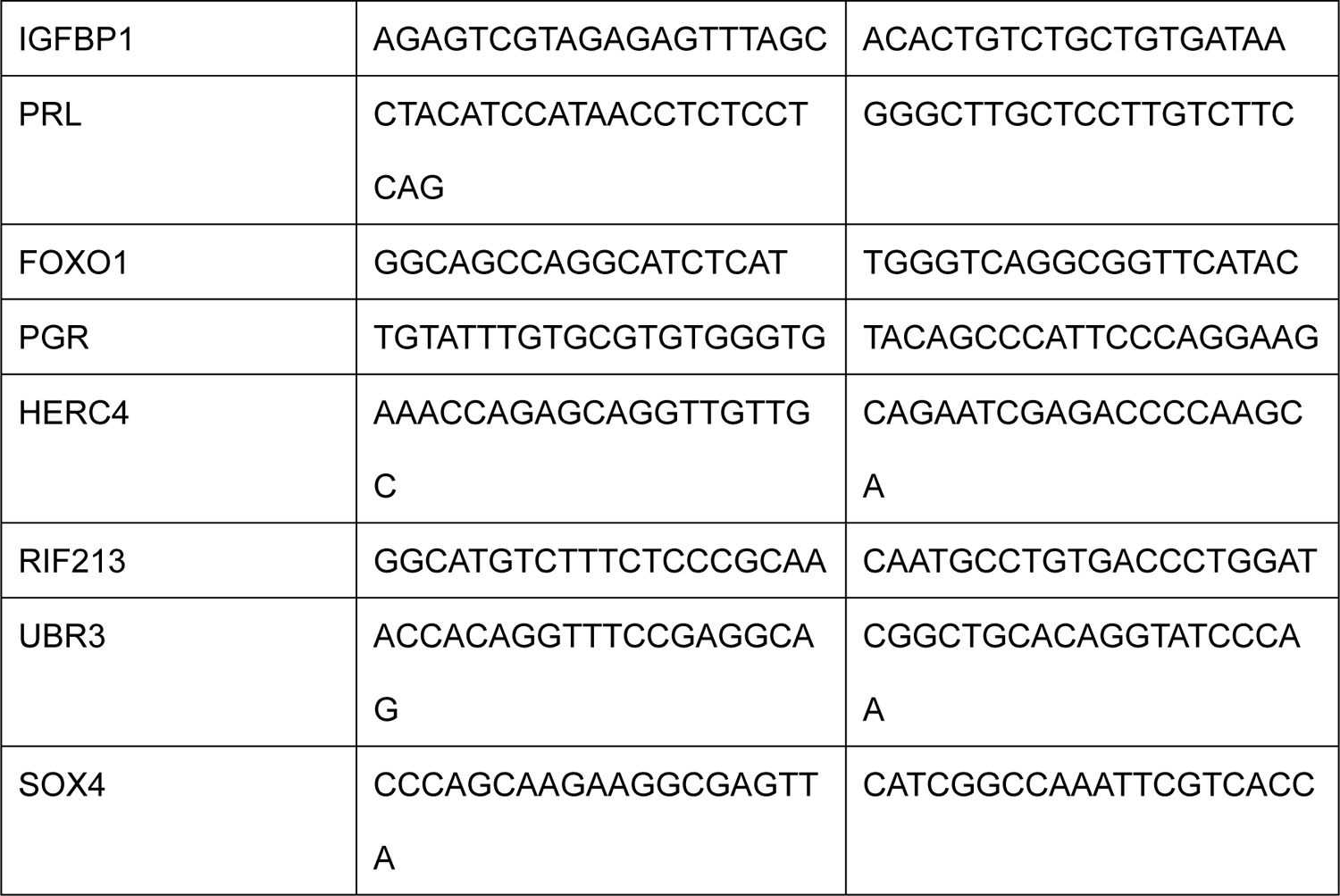
The sequence of oligonucleotide.

**Supplementary Table 1.** Detailed information of participants in this study

|  | Control(n=12) | EMS-RIF(n=12) | P |
| --- | --- | --- | --- |
| Age (years) | 32.62±3.50 | 32.65±3.43 | >0.05 |
| BMI (kg/m<sup>2</sup>) | 22.01±2.62 | 21.99±3.21 | >0.05 |
| Basal FSH (IU/l) | 7.00±2.23 | 7.05±3.50 | >0.05 |
| Duration of infertility (years) | 0 | 5.61±3.35 | >0.05 |
| No. of ET failures | 0 | 5.50±2.00 | >0.05 |
| Endometrial thickness (mm) | 11.29±3.01 | 10.89±2.65 | >0.05 |

